# Completed genomes from *Variovorax* provide insight into genome diversification through horizontal gene transfer

**DOI:** 10.1101/2025.03.19.644167

**Authors:** Christopher J. Ne Ville, Paul M. Orwin

## Abstract

Approximately 10% of all bacterial genomes sequenced thus far contain a secondary replicon. This property of bacterial populations vastly increases genomic diversity within phylogenetically narrow groups. Members of the genus *Variovorax* have extensive heterogeneity in genome architecture, including sequenced isolates containing plasmids, megaplasmids, and chromids. Many of the *Variovorax* genomes in the NCBI database were generated using only short-read data and were assembled to the permanent draft stage. We acquired a set of these isolates and used the MinION long-read sequencing device to generate additional data to allow hybrid assemblies of these genomes. Here we present the finished assemblies of 15 *Variovorax* isolates from diverse ecosystems that were previously only available as permanent drafts. When added to the previously published *Variovorax* assemblies for EPS, CSUSB, and VAI-C and those published by other groups, we found significant diversity in genome architecture. We found that there are plasmids, megaplasmids, and chromids that are distinguishable using Guanine-Cytosine (G+C content) content as a signal. We identified a plasmid integration event in NFACC27 and identify potential evolutionary relationships in the second replicons based on ParB homology as well as ANI. The evidence suggests that *Variovorax* is like its sister taxon *Burkholderia,* highly capable of acquiring and maintaining stable secondary replicons. The plasticity of these architectures and the mechanisms for maintenance remain a topic for future research.

## INTRODUCTION

Genome architecture in bacteria is now recognized as a major component of genetic and genomic variability [1–3]. Although the traditional model of a single circular chromosome and smaller circular plasmids is the rule, the architecture of many genomes is more complex. Linear plasmids, megaplasmids, secondary chromosomes, chromids, and linear chromosomes have been identified in diverse species [3,4]. The important role of stable secondary replicons is well established, starting with the symbiosis megaplasmid of *Sinorhizobium meliloti* [5]. Sequencing and assembly projects across the diversity of bacteria have shown that about 10% of all bacteria contain large, stable secondary replicons [4].

The four established types of circular secondary replicons are plasmids, megaplasmids, chromids, and secondary chromosomes. Plasmids are smaller elements with highly variable carriage across the taxon, and no essential genes present [4]. A megaplasmid is a large low copy number plasmid, with a rough cutoff of ∼400 kbp delineating a megaplasmid [4]. A chromid is a subdivision defined by the presence of at least one core gene, meaning the loss of this replicon would result in cell death [6]. The term chromid is a reflection of the intermediate status of this element between a plasmid and chromosome [4]. Several of the replicons termed secondary chromosomes of *Burkholderia* are best characterized as chromids [7], as are the previously identified second chromosomes of *Variovorax paradoxus* B4 [8], S110 [9], and VAI-C [10]. A secondary chromosome shares the same functional role as a chromid but is defined by its unique evolutionary origin, as the result of a fracture of the primary chromosome into two independently maintained replicons [4].

The plasmid invasion hypothesis [11–13] is a model for chromid formation in which the plasmid accumulates DNA from the host chromosome through recombination events. These recombination events enlarge the plasmid and make it less likely to move via horizontal transfer. This also brings the genomic signatures of the plasmid more in line with the chromosome. These signatures, which include G+C%, dinucleotide relative abundance (DNRA), and codon usage, influence gene expression but are independent of gene content, and thus can be used as phylogenetic signals [6]. The convergence of these signatures between primary and secondary replicon is indicative of the process of “domestication” of the plasmid [4,14]. The movement of an essential gene from the chromosome to the plasmid also stabilizes the second replicon in the population [6].

Multipartite genomes are distributed across the domain Bacteria but there are many clear clusters within certain phylogenetic groups, including the Burkholderiales. *Variovorax* is a metabolically diverse, aerobic, ubiquitous gram negative betaproteobacterial genus within this clade that features diverse genome architectures [15]. The G+C content of their genomes is between 66.5% and 69.4% with optimal growth temperatures of around 30°C [16]. Within *Variovorax* are strains that grow autotrophically by oxidizing hydrogen gas as well as plant growth promoting bacteria [15]. Important enzymatic activities such as quorum quenching [17], bioremediation [18], and enantiomeric selective drug synthesis [19] are associated with this genus. Many strains in this genus have been isolated from the rhizospheres of various wild and crop plants, and enzymatic activities associated with plant growth promotion (PGP) have been frequently demonstrated [9,20]. Rhizosphere *Variovorax* use mechanisms of auxin synthesis and ethylene degradation to promote root development [21] and are key determinants of bacteria-plant communication networks in complex model communities [21].

Multiple members of this genus have taken up and maintained large pieces of DNA, creating extensive genomic heterogeneity within the clade [9,22]. As of December 2023, 25 previously fully assembled genomes from *Variovorax* are in the NCBI genomes database along with many other “permanent draft” genomes. Some of these were published by our group previously [9,10,23,24], or by other groups identifying bioremediation [22,25]. The list of all the previously finished genomes along with those presented here (**Supplemental Table 1**) shows the heterogeneity of the architecture. Twelve of the previously published genomes have single circular chromosomes, while in the 13 others we see plasmids, megaplasmids, and chromids present.

Here we present the assembled genomes of 15 *Variovorax* isolates from a variety of sources, including the original type strain of *Variovorax paradoxus*. Strains from model rhizosphere communities, wild plant rhizospheres, freshwater lakes, and enrichment experiments are presented and compared to previously assembled genomes. We used Average nucleotide identity (ANI) and marker gene based phylogeny to examine the relationships between these genomes. We observed evidence of a megaplasmid integration event, and provide evidence for the classification of some of these elements as chromids.

## METHODS

### Variovorax isolates

Nine *Variovorax* isolates were acquired from Jeff Dangl at the University of North Carolina at Chapel Hill, one from Jared Leadbetter at Caltech, one from Kostas Konstaninidis at Georgia Tech, and four from the Noble Research Center (**Table 1**). Previously published data from strains with finished assemblies in the NCBI database were also used (**Supplemental Table 1**) for *in silico* comparisons.

**Table 1.** Genomes of *Variovorax* strains completed in this study.

| Isolate name | Genbank<br>Accession number | Short Read Data<br>Source |
| --- | --- | --- |
| *V. paradoxus* 4MFCol3.1 | GCA_048310315.1 | Dangl Lab |
| *V. paradoxus* 110B | GCA_048310285.1 | Dangl Lab |
| *Variovorax* sp. 160MFSha2.1 | GCA_048305075.1 | Dangl Lab |
| *Variovorax* sp. 278MFTsu5.1 | GCA_048305045.1 | Dangl Lab |
| *V. paradoxus* 295MFChir4.1 | GCA_048307675.1 | Dangl Lab |
| *V. paradoxus* 349MFTsu5.1 | GCA_048307605.1 | Dangl Lab |
| *Variovorax* sp. 350MFTsu5.1 | GCA_048302515.1 | Dangl Lab |
| *V. paradoxus* 369MFTsu5.1 | GCA_048307575.1 | Dangl Lab |
| *Variovorax* sp. 375MFSha3.1 | GCA_048302465.1 | Dangl Lab |
| *V. paradoxus* ATCC17713* | GCA_048305125.1 | This work |
| *Variovorax* sp. KK3 | GCA_048302415.1 | SRA |
| *Variovorax* sp. NFACC26 | GCA_048301555.1 | SRA |
| *Variovorax* sp. NFACC27 | GCA_048301545.1 | SRA |
| *Variovorax* sp. NFACC28 | GCA_048301535.1 | SRA |
| *Variovorax* sp. NFACC29 | GCA_048301405.1 | SRA |

\*This strain is also referred to as NBRC15149

### Genomic DNA isolation and library preparation

The genomic DNA and culture conditions were used as described in previous publications [10]. The protocol is based on the bacterial genomic CTAB DNA isolation protocol suggested by the Department of Energy’s (DoE) Joint Genome Institute (JGI) and Loman/Quick protocol [26,27]. An isolated colony for each isolate freshly streaked from a stock culture was used to inoculate 5 g/L YE broth. Cultures were grown into log phase at 30°C with shaking as determined by optical density (OD_600_=∼0.4-0.7). Cells were centrifuged at 17,000 x g for 10 minutes and the culture fluid was decanted and discarded. The cell pellets were stored at −80 °C prior to DNA extraction. Cell pellets were thawed on ice and resuspended in TE (10mM Tris, 1mM EDTA, pH 8). The JGI-DOE bacterial genomic DNA isolation using CTAB protocol [27] was followed with the following modifications. After incubation for 30 minutes at 37°C, 20 µL RNAse (Promega) was added to the solution. To denature the exopolysaccharide produced by many *Variovorax* isolates, the samples were incubated at 57°C overnight after addition of 10% SDS and Proteinase K. The lysates were extracted twice with phenol:chloroform:isoamyl alcohol and twice with chloroform:isoamyl alcohol. The aqueous layer was then removed and precipitated in ice-cold of absolute ethanol with 0.3 M ammonium acetate (final concentration). The sample was stored at least overnight at −20°C to allow for DNA precipitation. The precipitated DNA was removed using a sterilized glass capillary hook and submerged in 70% ethanol until it constricted around the capillary tube into a white pellet. The pellet was scraped into a 1.7mL centrifuge tube along with 1 mL of freshly made 70% ethanol followed by centrifugation at 4°C at 17,000 x g for 10 minutes. The supernatant was removed, and the tube was dried in a heated DNA speed-vac set to medium speed for 10 minutes. The isolated genomic DNA was resuspended in 100 µL of 10mM Tris (pH 8.0) and heated at 37°C for 10 minutes to dissolve the pellet fully. The concentration and purity of the DNA before library preparation were determined using a Nanodrop spectrophotometer (Thermo Fisher). A 26-gauge needle was used to shear the DNA before Nanopore library preparation. Nanopore libraries were constructed using the Rapid barcoding kit (SBK-004). In the instances where short-read data was also needed, the same genomic DNA preparation was used to generate a 250-300 bp library with Nextera DNA Flex LPK kit. Nanopore sequencing was performed in the MinION device in a R9.4.1 (Min-106) flow cell, and short read sequencing was performed using the iSEQ (illumina).

### Genome Assembly Pipeline

The raw Nanopore data was basecalled using Guppy (v2.3.1) configuration r9.4.1 Flipflop v1.1.0 or Flappie (later High-Accuracy Basecalling [HAC]), running on standard configuration [28]. MinION reads were demultiplexed in Deepbinner [29] and Guppy with the agreed upon reads adapters removed with Porechop (v0.2.4) [30]. Quality control of the long reads was done by running the reads in Filtlong (v0.2.0) [31] using the Illumina data as an external reference to throw out low identity reads and barcodes that slipped through the demultiplexing process. For the Illumina data FastQC (v0.11.8) was used for quality assessment of this data [32]. Trimming was performed in Trimmomatic (0.38.0) [33], applying the IlluminaCLIP procedure to remove any remaining sequence adaptors/barcodes. Also in Trimmomatic, the first 20 nucleotides were cropped off each read (HEADCROP), and the last five were also removed (TRAILING), leaving a mean length distribution of 126 bp. Short read data from the Dangl lab strains (**Table 1**) was provided by Dangl Lab at the University of North Carolina at Chapel Hill, while short read data pertaining to KK3 and the Noble group strains were obtained from public datasets on the JGI website (www.img.jgi.doe.gov).

Assemblies were created using a hybrid approach in Unicycler (v0.4.8.0) [30] utilizing default parameters without references on the North America Galaxy Hub (http://usegalaxy.org) [34]. For any assembly which did not circularize in Unicycler, Nanopore reads were aligned to regions of discontinuity using Minimap2 (v2.20) [35] to create a .sam file. Samtools (v1.12) [36] was used to sort/index the file into a .bam file which could be viewed in IGV (v2.10.0) [37]. Reads which aligned to the area of discontinuity were extracted to a fastq file. This fastq file was then used in Bandage [38] to create a BLAST bin to confirm they spanned the gap. This bin was used to create a consensus read in ALFRED (v.0.2.3) [39]. Initial annotation was performed by uploading the assembly to Rast.org and using the RASTtk pipeline (http://rast.nmpdr.org) [40]. The assemblies along with the raw data were then submitted to the NCBI database which uses the PGAP pipeline prior to making the assemblies available to the public.

### Core Gene Identification

Core genes in the secondary replicon were identified by taking Betaproteobacteria 203 gene .hmm file from GtoTree (v.1.5.44) [41] and using it as a base for a HMMER3 (v.3.3.2) [42] search.

### Replicon characteristics

G+C content was determined from database values for previously published replicons or the G+C value calculated for the Kbase annotation was used. Sizes of individual replicons were obtained similarly. Secondary replicons >1 Mb are identified as chromids, replicons < 300 kB were identified as plasmids, and intermediate length replicons are termed megaplasmids. Average Nucleotide Identity data for pairwise comparisons was generated using ANI (http://enve-omics.ce.gatech.edu/ani/) [43].

### Dotplot

The dotplot was constructed using the dotplot tool in RAST [40] to align the genome of *Variovorax* NFACC27 with the megaplasmid assembled from NFACC29.

### Marker Based Phylogenomic Tree

A genome assembly set was created in Kbase including all of the finished annotated genomes from the NCBI database as of December 2023, and the finished genomes in this work. The marker based phylogenetic tree was generated using SpeciesTree v2.2 [44] tool in Kbase (www.kbase.us). The tree was visualized in TreeViewer [45] and excess neighboring genomes were pruned or collapsed.

### ParB based tree

The JACKHMMer [42] tool was employed to search a dataset constructed from the 40 secondary replicons in the finished genome dataset (see **Supplementary Data**). The dataset was built by concatenating the amino acid FASTA files (.faa) from Prokka [46] annotations of the secondary replicons. The ParB sequence from the VAI-C chromid annotation was used at the search query with maximum of five iterations. The same search was conducted using the ParB sequence from 4MFCol3.1 with no difference in outcome. The list of hits was filtered with an E-value cutoff of 1×10^−10^ resulting in seventy retained sequences. These sequences were extracted from the dataset and aligned using MUSCLE [47] along with an outgroup ParB from *Xanthomonas campestris*. A tree based on that alignment was generated in IQ-tree [48] and the downloaded Newick tree was then visualized in R [49] using a custom script (**Supplementary Data).** All of the analysis prior to visualization was conducted using the Galaxy Europe server (www.usegalaxy.eu) [34].

### VAI-C chromid analysis

The analysis of the VAI-C chromid was undertaken using the ProkSee platform [50]. The Alien Hunter algorithm [51] was used to identify horizontal gene transfer regions within the chromid.

### Pangenome

The pangenome comparisons were made using *V. paradoxus* NBRC15149 (also known as ATCC17713, the type strain for *V. paradoxus*) as the basis using the ComputePangenome tool (v0.0.7) in Kbase [52]. The Kbase Pangenome Circle Plot [53] tool (v1.2.0) was then used to visualize the comparisons.

### Data Availability Statement

Previously published data can be found in NCBI using accession numbers already reported. Newly available data can be found in BioProject PRJNA1112457 and newly released assemblies are available (BioSample Accession SAMN41427624-38). The complete genome assembly accession numbers are found in **Table 1**. The individual genome pieces (chromosomes, chromids, plasmids) are available with accession numbers CP157610-CP157637). FastQ files for the data collected in this work are available in the same BioProject (SRA accession PRJNA1112457). Raw .fast5 files are available upon request.

## RESULTS AND DISCUSSION

### Replicon Identification

For the 15 completed assemblies, the fasta files for each individual replicon was extracted for further analysis. A summary of replicon sizes and GC content of all known secondary replicons is presented in **Supplemental Table 1**. The core genes identified on the newly assembled secondary replicons are listed in **Supplemental Table 2**. The second replicon in *V. paradoxus* 110B has G+C content and size characteristics similar to the chromids in other closely related strains, but no core genes were identified on the secondary replicon. None of the core genes found on the secondary replicons were missing from the primary chromosome.

### Variovorax whole genome phylogeny

A marker-based tree (**Figure 1**) generated using that shows the genus *Variovorax* is a phylogenetically consistent (monophyletic) group. However the species delineations are not as clear. This is also reflected in the current GDTB delineations of *Variovorax,* and indicates that the species naming in the genus needs revisiting.

**Figure 1.**
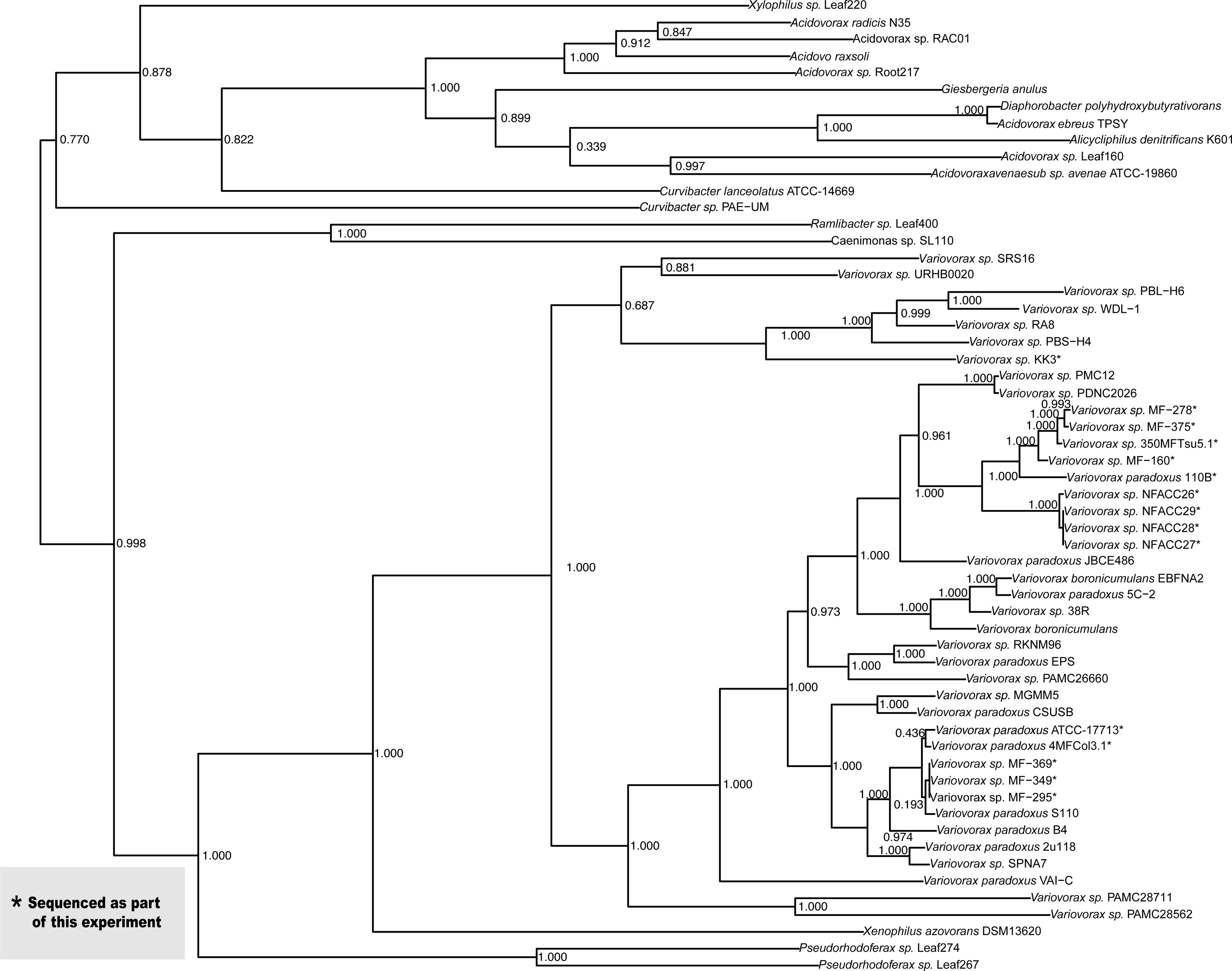
Marker Based Phylogenetic Tree Built using Maximum Likelihood. Strains with complete genomes were placed into a species tree along with neighboring organisms. The tree was built using all conserved markers from betaproteobacteria. Newly reported genomes from this work are starred.

### Secondary Replicon Types

In the set of finished genomes, the sizes of secondary replicons range from 20kb to 2.4Mb, and as described. Of the newly identified second replicons reported in this work, we identified seven putative chromids, and nine putative plasmids (size range 260-570kb). **G+C Content**. The G+C contents of plasmids and megaplasmids are more broadly distributed than chromids and primary chromosomes. (**Figure 2**). In *Variovorax,* the G+C contents of the chromosomes were always higher than those of the secondary replicons (**Figure 2)**. This is consistent with larger studies showing that G+C contents of plasmids are are usually lower than the chromosomes, even in gammaproteobacteria and spirochaetes which have overall very low G+C contents [6]. The consistency of this relationship suggests a possible mechanistic reason for this pattern.

**Figure 2.**
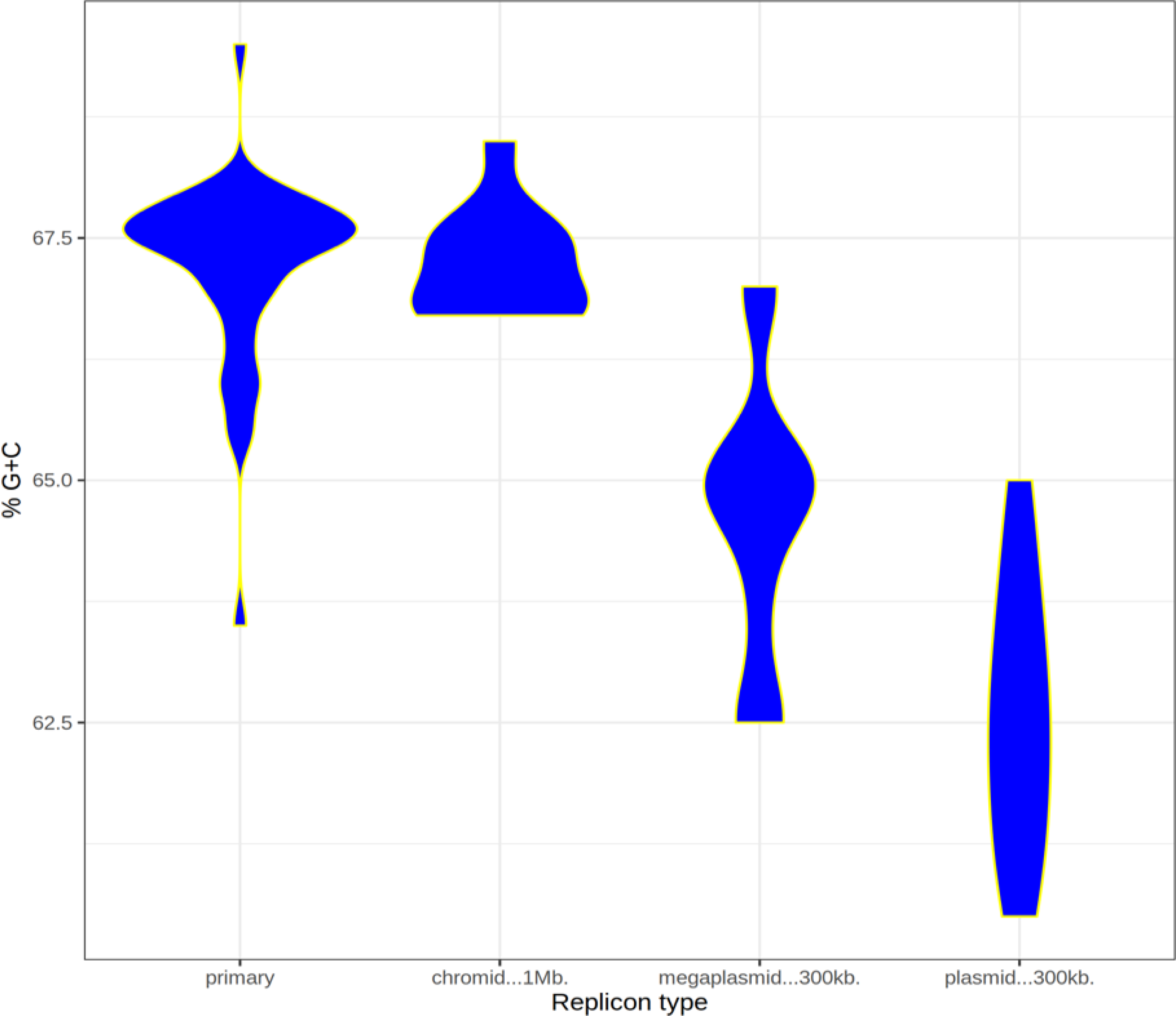
Violin plot of G+C content of different types of replicons showing that mobile elements and putative MGE derived elements have lower G+C than primary chromosomes in *Variovorax*.

### VAI-C has an unusually large second replicon

The genome of VAI-C was published previously, and the second replicon was notably larger than in other complete assemblies. A G+C content plot of the second replicon (**Supplemental Figure 1**) identifies a large section of this assembly with a notably lower G+C content than the rest of the element. Using the Alien Hunter algorithm [51] we examined this replicon and the results indicate a large insertion event, or possibly multiple insertion events into this region of the plasmid (**Supplemental Figure 1**). CheckM [54] analysis did not indicate any contamination of the assembly, and random gene checks using tBlastn [55] to compare predicted amino acid sequences with genomes routinely return genes from *Variovorax* isolates or other betaproteobacteria, frequently from plasmids.

### Identification of a megaplasmid integration event

*Variovorax sp.* NFACC27 is nearly identical to NFACC26,28, and 29, but does not contain an obvious secondary replicon. Analysis of the finished genomes provides evidence for the recent integration of the megaplasmid found in these other strains. The integration site was identified by using closely related organism’s (NFACC26, NFACC27, NFACC29) megaplasmids to BLASTn search the chromosome. To verify the integration site, both Nanopore and Illumina reads were aligned to see if there were reads that spanned from the chromosome, over the integration site and to the megaplasmid. There were short and long reads that spanned from the NFACC27 chromosome onto the region that aligns with the megaplasmid, suggesting that this was not an artifact of sequencing or an assembly error. A dot plot comparing NFACC27 and NFACC29 (**Figure 3)** shows the presence of the megaplasmid at around 4Mbp. Based on the level of sequence similarity between these strains it seems likely that this was a recent event.

**Figure 3.**
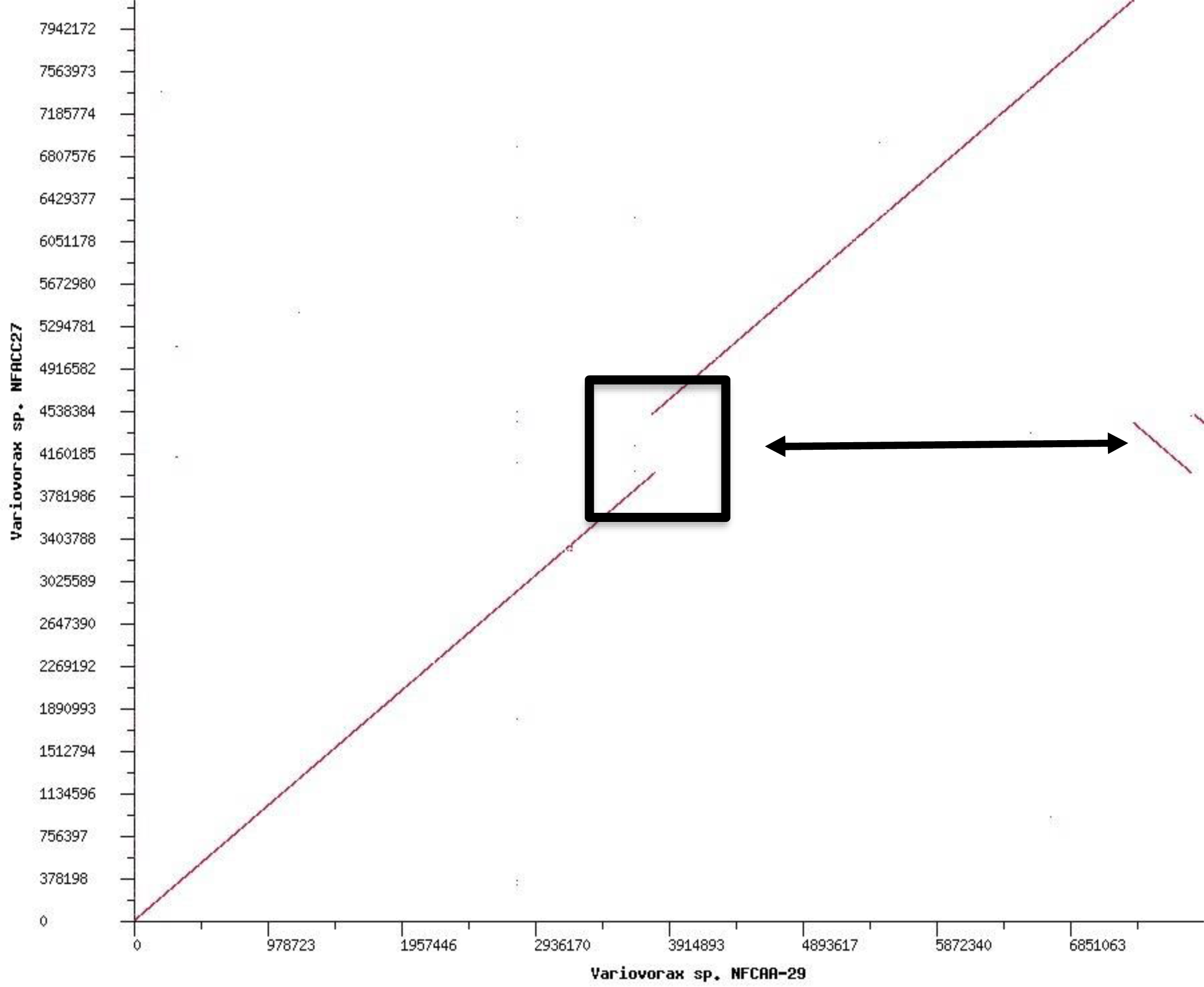
NFACC27 Megaplasmid Integration Event. Dotplot of NFACC29 megaplasmid against the NFACC27 genome illustrating the integration of the NFACC27 megaplasmid at approximately 4 Mbp.

### Average Nucleotide Identity

We used pairwise ANI across of the completed Variovorax genomes to evaluate relatedness (**Supplemental Data Figure 2)**. There are several organisms classified as *V. paradoxus* that fall below the species delineation cutoff at 95-96% when compared to other organisms classified as being in this species (EPS, VAI-C, B4). The cluster map built from this data (**Figure 4**) further reinforces the idea that species delineation within the genus is murky. **Figure 4** shows that many of the organisms which were previously named *V. paradoxus* do not cluster well with the others of their species. We further observe from this data that the organisms carrying similar replicon types do cluster together. The overall genome cluster map aligns well with the marker-based tree (**Figure 1**). A similar ANI approach was used to cluster the secondary replicons (**Supplemental Figure 2)** resulting in the diagram in **Figure 5**. The topography of this cluster map suggests strongly that independent replicon invasions have occurred repeatedly, not just with the smaller plasmids in strains such as WDL-1, SRS-16, RA8, PBL-H4, PBL-H6, and PBL-E5, but also with the larger megaplasmids and chromids. At least 5 distinct groups of secondary replicons are clearly identified in the NFACC strains, the ATCC type strain and its close relatives, the VAI-C replicons, the WDL-1 and SRS-16 plasmids, and the KK3 plasmid. The cultivation of multiple strains from a linuron contamination site in Belgium (**Supplemental Table 1**) is the straightforward explanation for the plasmids shared among the strains from that site, but the association of some of those plasmids with geographically distant strains suggests deeper layers of HGT earlier in the evolution of these strains.

**Figure 4.**
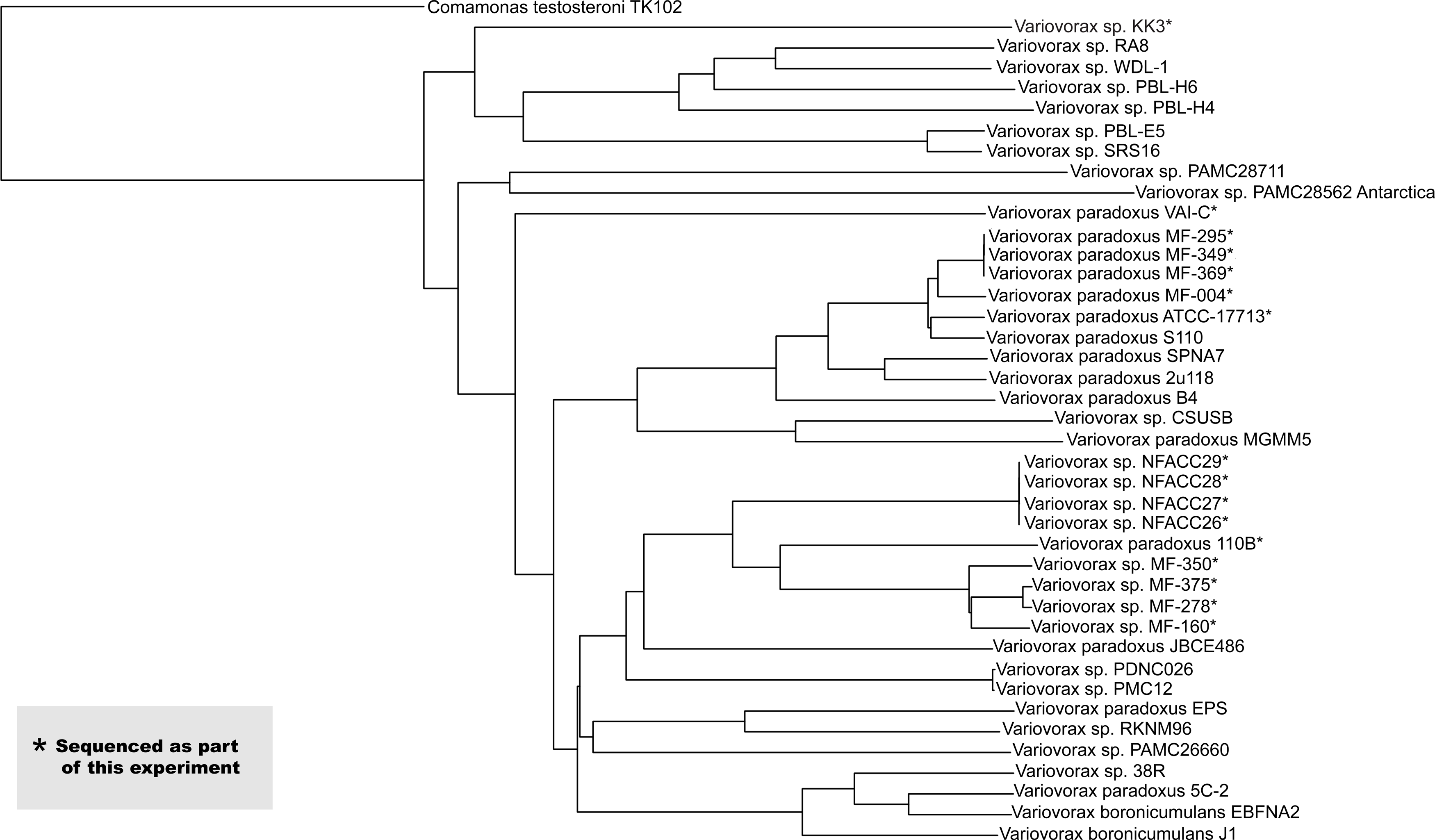
ANI cluster diagram of all complete *Variovorax* genomes. This cluster map was generated from the genomes in **Table 2** and including *Comamonas testeroni* TK102 as an outgroup. Starred genomes were sequenced as part this experiment.

**Figure 5.**
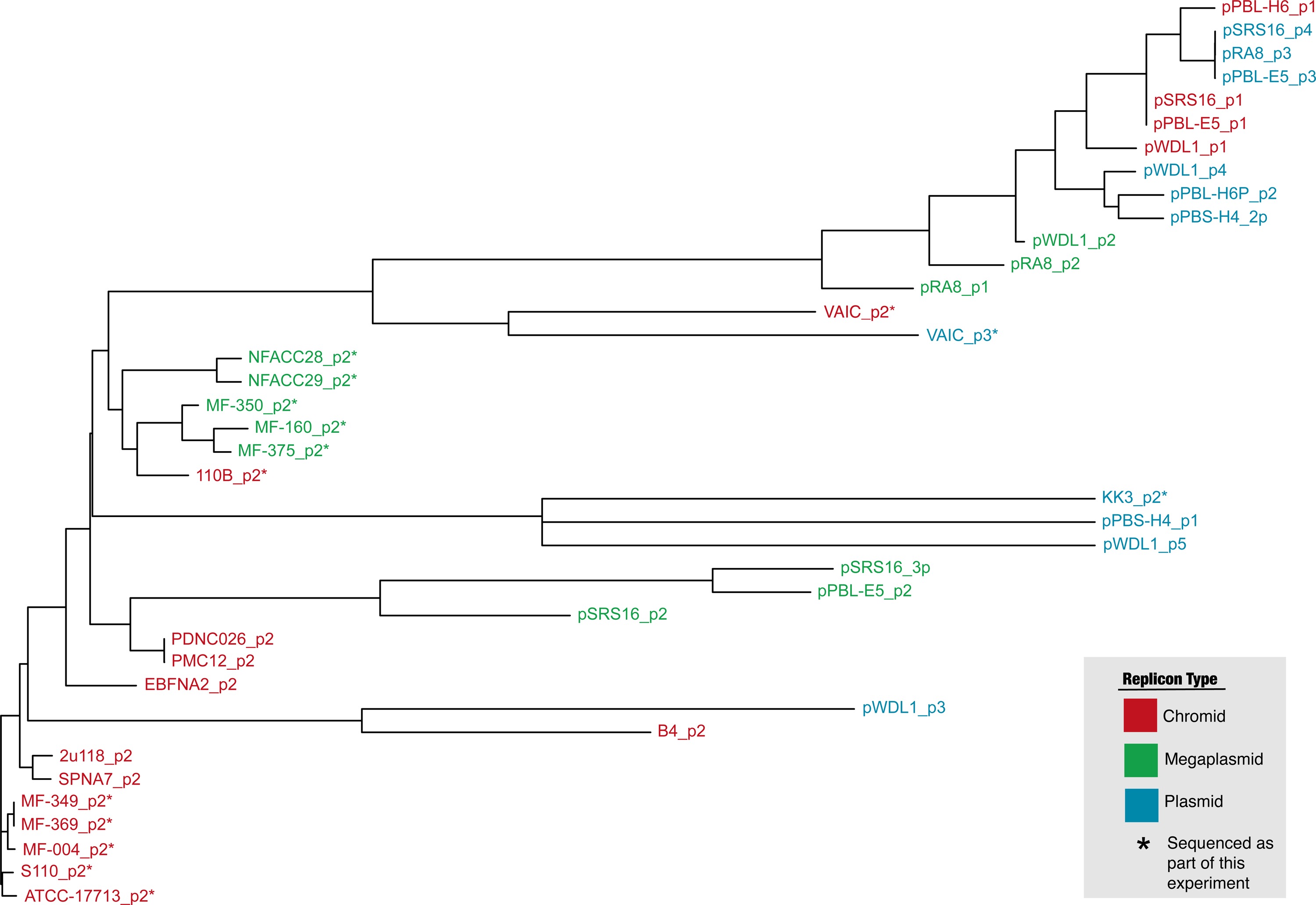
ANI cluster diagram of second replicons. ANI was performed using only the secondary replicons. The colors reflect the different replicon types as described in the introduction, and the starred replicons are the ones reported for the first time in this work.

### Marker based plasmid analysis

A ParB based tree on the secondary replicons (**Figure 6**) shows that shows that the genomic architectural diversity observed is likely from multiple different HGT events and not a single HGT event and diversification. This tree also shows that a diverse set of plasmid maintenance systems are present in the genus. The relatedness of certain ParB orthologues present in geographically distant isolates suggests pervasive horizontal transfer of these elements, and the polyphyletic nature of the ParB tree indicates that these plasmids are able to move widely in the betaproteobacteria.

**Figure 6.**
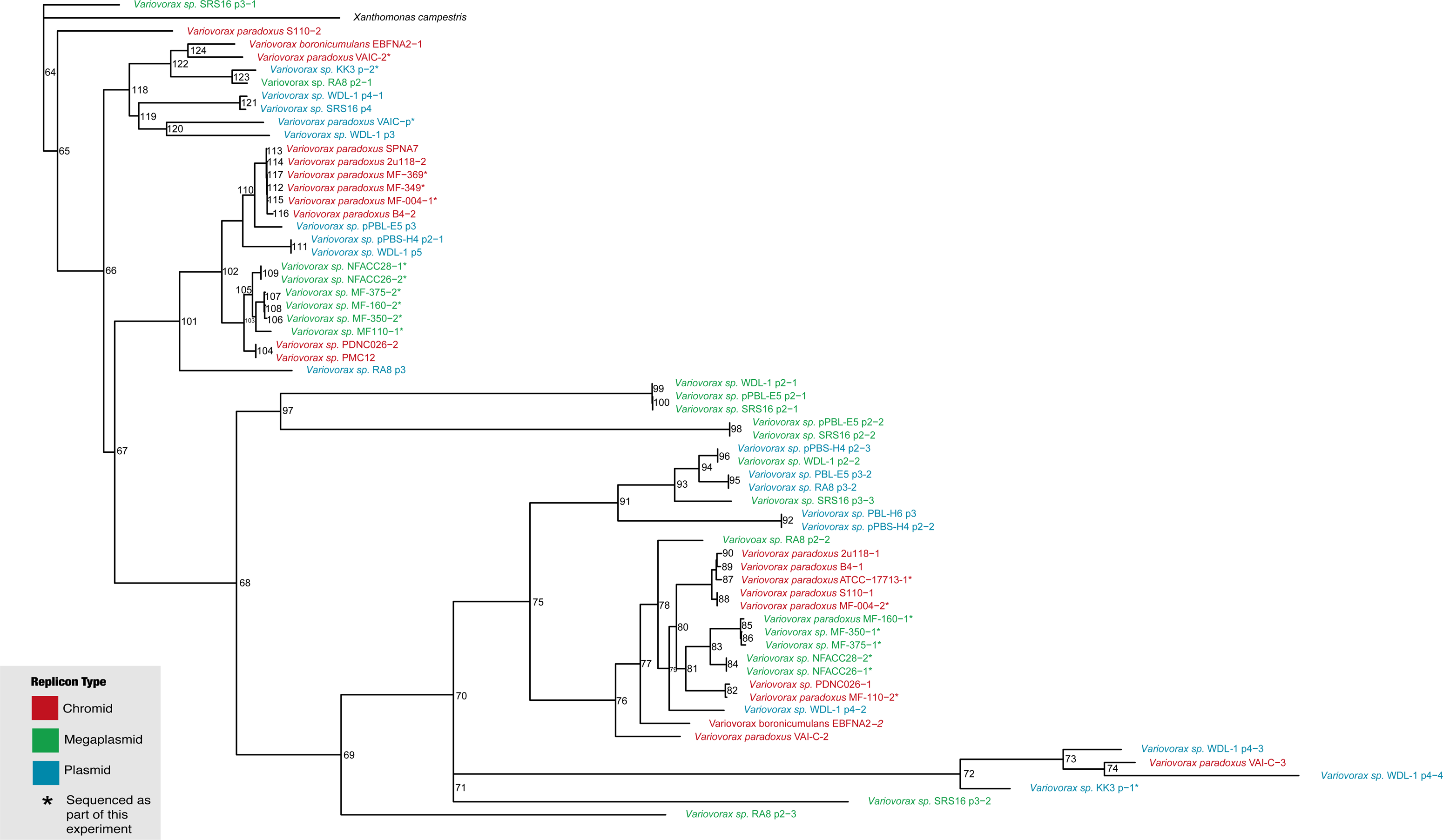
ParB based phylogeny of secondary elements. The single gene maximum likelihood tree built from an alignment of ParB amino acid sequences has branching patterns similar to the whole genome phylogeny (Fig 1). Duplicate ParB orthologs present in individual replicons are possibly indicative of composite plasmids formed by integration or fusion events. Sequences derived from replicons sequenced for this work are starred. Bootstrap values are shown at each branchpoint.

Support for the accretion model of large secondary replicon evolution [56] is also seen in this ParB based tree. Several aspect of the tree support this model, including the presence of multiple ParB homologs in a single replicon, and the presence of close ParB homologs in all 3 types of replicon. The most parsimonious explanation for this pattern of relatedness is that larger elements form by the fusion of smaller elements along with the movement of DNA from the chromosome to the secondary element.

Focusing on the VAI-C replicons and comparing the ANI and ParB trees suggests an intricate evolutionary history. The multiple ParB genes in the larger chromid are spread throughout the tree and are closely related to a diverse set of loci in different plasmids, megaplasmids, and other chromids. The ANI results, however, cluster for the two VAI-C replicons are more closely than any others. The most parsimonious explanation for this pattern is the fusion of multiple plasmids or megaplasmids to form the larger chromid, followed by duplication and recombination between the elements once stably maintained. The megaplasmid fusion hypothesis is further supported by the observed G+C and HGT pattern described earlier (**Supplemental Figure 1).** This evolutionary path is only evident by combining marker based and ANI based approaches. Additionally, the presence of multiple ParB orthologs in individual replicons (up to 4 in the WDL1p4) suggest plasmid fusion events in the evolutionary history of these plasmids. The segregation of these ParB variants into clusters in the tree indicates that the event occurred before the plasmids diverged. Overall this data suggests that recombination between the elements of the genome, including the chromosome as seen in NFACC27 and the plasmids present in many strains, may be a common event within the genus *Variovorax*.

### Pangenomes

The pangenome plots in **Figure 7** show a strong primary replicon bias in conserved genes, as predicted by the ANI. The primary replicons of the newly finished genomes show large regions of highly conserved genes across the breadth of the genus (**Figure 7A)**, while the secondary replicons share few if any genes across all replicons (**Figure7B**). We find one cluster of closely related chromids (**Figure 7B**), corresponding to the clustered secondary replicons in **Figure 5** and **6**.

**Figure 7.**
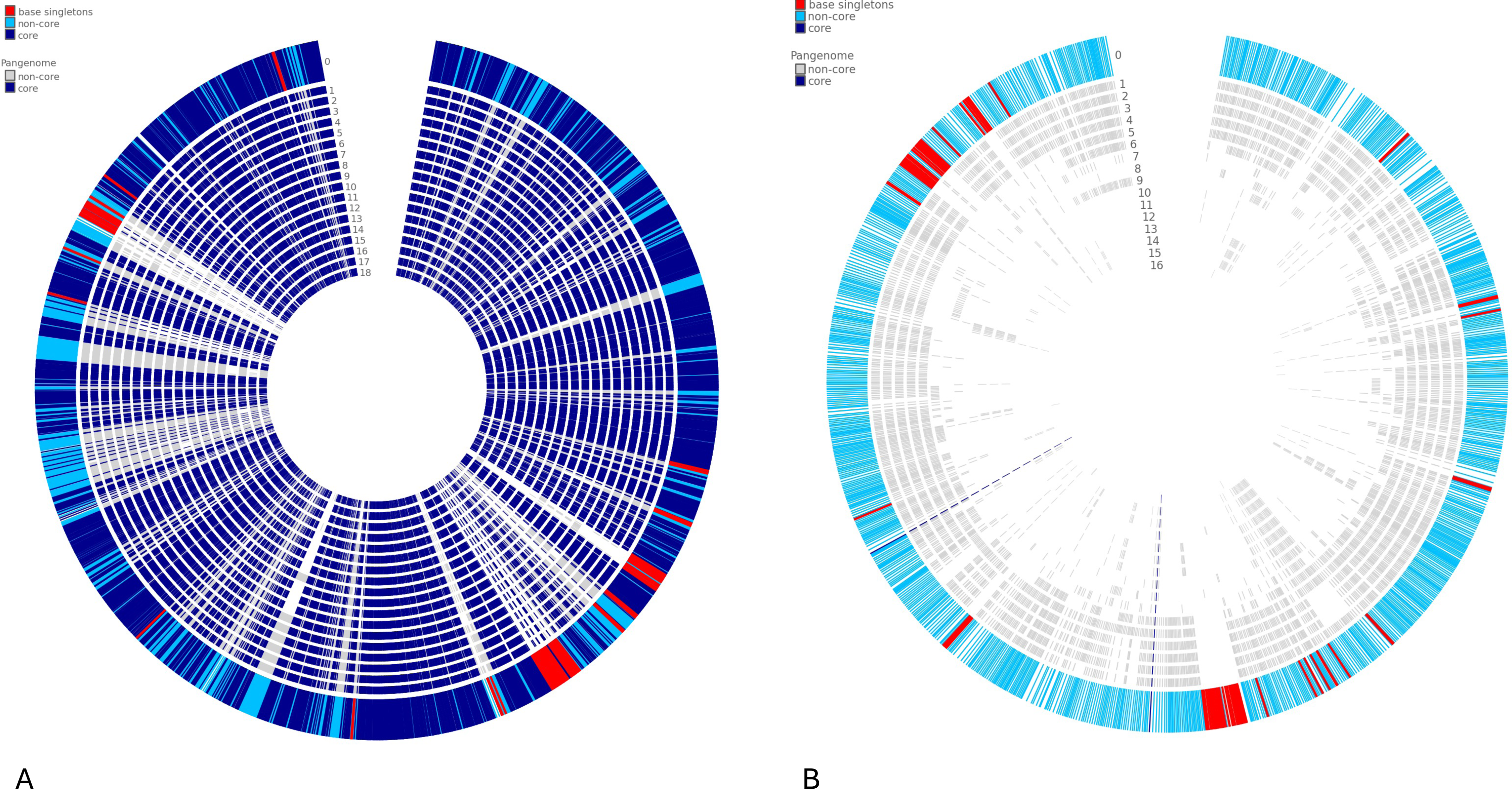
Pangenome circle plot of newly assembled primary replicons (A) and secondary replicons (B). Pangenomes both use the replicons from the ATCC17713 type strain of *Variovorax paradoxus* as the base. A cluster of closely related second replicons is seen in the outer rings of (B).

## CONCLUSION

The finished genome assemblies we report here help us build a clearer picture of the phylogenetic relationships in the genus Variovorax. The identification of multiple variable secondary replicons, as well as the identification of a replicon integration event, illustrate the importance of completing genome assemblies and not just leave them as permanent draft assemblies. With the 15 assembled *Variovorax* genomes, we report here along with those previously published or submitted from other research groups we can see there is extensive heterogeneity. This pattern of genome architecture is consistent with plasmid acquisition and expansion over evolutionary time. Marker based analysis using the ParB plasmid replication protein loci indicates further evolutionary complexity. Many of the plasmids contain multiple genes encoding ParB orthologs, indicating that plasmids, megaplasmids, and chromids form and grow by fusion of previously independent elements. The complexity of the marker based phylogeny suggests that these events are frequent. The G+C content analysis supports the hypothesis that uptake of a plasmid is followed by DNA transfer and mutation that leads to regression of the DNA content to match the composition of the chromosome. The observation of a megaplasmid integration event, along with the observation that multiple isolates within the group have singular chromosomes, suggests that the genome architecture may be fluid. The regulation of these processes remains an open question to be studied further, but there may be diverse environmental and genetic factors that lead to the differential propensity to acquire and maintain secondary replicons.

## ACKNOWLEDGEMENTS

This work was funded by grants provided by California State University, San Bernardino to PMO and CJN. Strains for sequencing were generously provided by Jared R Leadbetter, Jeff Dangl, and the Noble Research Institute.

## REFERENCES

1. Dame RT, Rashid F-ZM, Grainger DC. Chromosome organization in bacteria: mechanistic insights into genome structure and function. Nat Rev Genet. 2020;21: 227–242.

2. Kirchberger PC, Schmidt ML, Ochman H. The Ingenuity of Bacterial Genomes. Annu Rev Microbiol. 2020;74: 815–834.

3. Arnold BJ, Huang I-T, Hanage W. Horizontal gene transfer and adaptive evolution in bacteria. Nat Rev Microbiol. 2021;20: 206–218.

4. diCenzo GC, Finan TM. The Divided Bacterial Genome: Structure, Function, and Evolution. Microbiol Mol Biol Rev. 2017;81. doi:10.1128/MMBR.00019-17

5. Finan TM, Weidner S, Wong K, Buhrmester J, Chain P, Vorhölter FJ, et al. The complete sequence of the 1,683-kb pSymB megaplasmid from the N 2 - fixing endosymbiont Sinorhizobium meliloti. Proc Natl Acad Sci USA. 2001;98: 9889–9894.

6. Harrison PW, Lower RPJ, Kim NKD, Young JPW. Introducing the bacterial “chromid”: not a chromosome, not a plasmid. Trends Microbiol. 2010;18: 141–148.

7. Bochkareva OO, Moroz EV, Davydov II, Gelfand MS. Genome rearrangements and selection in multi-chromosome bacteria Burkholderia spp. BMC Genomics. 2018;19: 965.

8. Brandt U, Hiessl S, Schuldes J, Thürmer A, Wübbeler JH, Daniel R, et al. Genome-guided insights into the versatile metabolic capabilities of the mercaptosuccinate-utilizing β -proteobacterium V ariovorax paradoxus strain B 4. Environmental Microbiology. 2014;16: 3370–3386.

9. Han J-I, Choi H-K, Lee S-W, Orwin PM, Kim J, LaRoe SL, et al. Complete Genome Sequence of the Metabolically Versatile Plant Growth-Promoting Endophyte Variovorax paradoxus S110. J Bacteriol. 2011;193: 1183–1190.

10. Ne Ville CJ, Leadbetter JR, Orwin PM. Hybrid Assembly of the Quorum-Quenching Isolate Variovorax paradoxus VAI-C Genome Sequence. Thrash JC, editor. Microbiol Resour Announc. 2021;10. doi:10.1128/MRA.00265-21

11. Egan ES, Fogel MA, Waldor MK. Divided genomes: negotiating the cell cycle in prokaryotes with multiple chromosomes: Multiple chromosomes in prokaryotes. Mol Microbiol. 2005;56: 1129–1138.

12. Chain PSG, Denef VJ, Konstantinidis KT, Vergez LM, Agulló L, Reyes VL, et al. Burkholderia xenovorans LB400 harbors a multi-replicon, 9.73-Mbp genome shaped for versatility. Proc Natl Acad Sci U S A. 2006;103: 15280–15287.

13. Choudhary M, Cho H, Bavishi A, Trahan C, Myagmarjav B-E. Evolution of Multipartite Genomes in Prokaryotes. Evolutionary Biology: Mechanisms and Trends. Berlin, Heidelberg: Springer Berlin Heidelberg; 2012. pp. 301–323.

14. Cooper VS, Vohr SH, Wrocklage SC, Hatcher PJ. Why genes evolve faster on secondary chromosomes in bacteria. PLoS Comput Biol. 2010;6: e1000732.

15. Satola B, Wübbeler JH, Steinbüchel A. Metabolic characteristics of the species Variovorax paradoxus. Appl Microbiol Biotechnol. 2013;97: 541–560.

16. Willems A, Mergaert J, Swings J. Variovorax. Bergey’s Manual of Systematics of Archaea and Bacteria. American Cancer Society; 2015. pp. 1–9.

17. Leadbetter JR, Greenberg EP. Metabolism of Acyl-Homoserine Lactone Quorum-Sensing Signals by Variovorax paradoxus. J Bacteriol. 2000;182: 6921–6926.

18. Benedek T, Szentgyörgyi F, Gergócs V, Menashe O, Gonzalez PAF, Probst AJ, et al. Potential of Variovorax paradoxus isolate BFB1_13 for bioremediation of BTEX contaminated sites. AMB Express. 2021;11: 126.

19. Krieg L, Ansorge-Schumacher MB, Kula M-R. Screening for Amidases: Isolation and Characterization of a Novel D-Amidase from Variovorax paradoxus. Adv Synth Catal. 2002;344: 965–973.

20. Garcia Teijeiro R, Belimov AA, Dodd IC. Microbial inoculum development for ameliorating crop drought stress: A case study of Variovorax paradoxus 5C-2. N Biotechnol. 2020;56: 103–113.

21. Finkel OM, Salas-González I, Castrillo G, Conway JM, Law TF, Teixeira PJPL, et al. A single bacterial genus maintains root growth in a complex microbiome. Nature. 2020;587: 103–108.

22. Carbajal-Rodríguez I, Stöveken N, Satola B, Wübbeler JH, Steinbüchel A. Aerobic Degradation of Mercaptosuccinate by the Gram-Negative Bacterium Variovorax paradoxus Strain B4. J Bacteriol. 2011;193: 527–539.

23. Han J-I, Spain JC, Leadbetter JR, Ovchinnikova G, Goodwin LA, Han CS, et al. Genome of the Root-Associated Plant Growth-Promoting Bacterium Variovorax paradoxus Strain EPS. Genome Announc. 2013;1. doi:10.1128/genomeA.00843-13

24. Ville CN, Enright D, Hernandez I, Dodsworth J, Orwin P. Complete Genome Sequences of Pseudomonas alkylphenolica Neo and Variovorax sp. Strain CSUSB, Obtained in Undergraduate Microbiology Courses Using a Hybrid Assembly Approach. Microbiol Resour Announc. 2020;9. doi:10.1128/MRA.01520-19

25. Öztürk B, Werner J, Meier-Kolthoff JP, Bunk B, Spröer C, Springael D. Comparative Genomics Suggests Mechanisms of Genetic Adaptation toward the Catabolism of the Phenylurea Herbicide Linuron in Variovorax. Genome Biol Evol. 2020;12: 827–841.

26. Quick J, Loman NJ. DNA Extraction Strategies for Nanopore Sequencing. Nanopore Sequencing. WORLD SCIENTIFIC; 2018. pp. 91–105.

27. Number: V. Bacterial genomic DNA isolation using CTAB. Available: https://jgi.doe.gov/wp-content/uploads/2014/02/JGI-Bacterial-DNA-isolation-CTAB-Protocol-2012.pdf

28. Wick RR, Judd LM, Holt KE. Performance of neural network basecalling tools for Oxford Nanopore sequencing. Genome Biol. 12/2019;20. doi:10.1186/s13059-019-1727-y

29. Wick RR, Judd LM, Holt KE. Deepbinner: Demultiplexing barcoded Oxford Nanopore reads with deep convolutional neural networks. PLoS Comput Biol. 2018;14: e1006583.

30. Wick RR, Judd LM, Gorrie CL, Holt KE. Completing bacterial genome assemblies with multiplex MinION sequencing. Microb Genom. 2017;3. doi:10.1099/mgen.0.000132

31. Wick RR. Filtlong https://github.com/rrwick. Filtlong.

32. Andrews S. FastQC: A quality control tool for high throughput sequence data. 2010. Available: https://www.bioinformatics.babraham.ac.uk/projects/fastqc/

33. Bolger AM, Lohse M, Usadel B. Trimmomatic: a flexible trimmer for Illumina sequence data. Bioinformatics. 2014;30: 2114–2120.

34. Afgan E, Baker D, Batut B, van den Beek M, Bouvier D, Čech M, et al. The Galaxy platform for accessible, reproducible and collaborative biomedical analyses: 2018 update. Nucleic Acids Res. 2018;46: W537–W544.

35. Li H. Minimap2: pairwise alignment for nucleotide sequences. Bioinformatics. 2018;34: 3094–3100.

36. Danecek P, Bonfield JK, Liddle J, Marshall J, Ohan V, Pollard MO, et al. Twelve years of SAMtools and BCFtools. Gigascience. 2021;10: giab008.

37. Thorvaldsdóttir H, Robinson JT, Mesirov JP. Integrative Genomics Viewer (IGV): high-performance genomics data visualization and exploration. Brief Bioinform. 2013;14: 178–192.

38. Wick RR, Schultz MB, Zobel J, Holt KE. Bandage: interactive visualization of de novo genome assemblies. Bioinformatics. 2015;31: 3350–3352.

39. Rausch T, Hsi-Yang Fritz M, Korbel JO, Benes V. Alfred: interactive multi-sample BAM alignment statistics, feature counting and feature annotation for long- and short-read sequencing. Bioinformatics. 2019;35: 2489–2491.

40. Brettin T, Davis JJ, Disz T, Edwards RA, Gerdes S, Olsen GJ, et al. RASTtk: a modular and extensible implementation of the RAST algorithm for building custom annotation pipelines and annotating batches of genomes. Sci Rep. 2015;5: 8365.

41. Lee MD. GToTree: a user-friendly workflow for phylogenomics. Bioinformatics. 2019;35: 4162–4164.

42. Eddy SR. Accelerated profile HMM searches. PLoS Comput Biol. 2011;7: e1002195.

43. Jain C, Rodriguez-R LM, Phillippy AM, Konstantinidis KT, Aluru S. High throughput ANI analysis of 90K prokaryotic genomes reveals clear species boundaries. Nat Commun. 2018;9: 1–8.

44. Price MN, Dehal PS, Arkin AP. FastTree 2--approximately maximum-likelihood trees for large alignments. PLoS One. 2010;5: e9490.

45. Bianchini G, Sánchez-Baracaldo P. TreeViewer: Flexible, modular software to visualise and manipulate phylogenetic trees. Ecol Evol. 2024;14: e10873.

46. Seemann T. Prokka: rapid prokaryotic genome annotation. Bioinformatics. 2014;30: 2068–2069.

47. Edgar RC. MUSCLE: multiple sequence alignment with high accuracy and high throughput. Nucleic Acids Res. 2004;32: 1792–1797.

48. Nguyen L-T, Schmidt HA, von Haeseler A, Minh BQ. IQ-TREE: a fast and effective stochastic algorithm for estimating maximum-likelihood phylogenies. Mol Biol Evol. 2015;32: 268–274.

49. Gentleman R, Carey V, Huber W, Irizarry R, Dudoit S. Bioinformatics and computational biology solutions using R and bioconductor. 2005th ed. Gentleman R, Carey V, Huber W, Irizarry R, Dudoit S, editors. Springer Science+Business Media; 2005. doi:10.1007/0-387-29362-0

50. Grant JR, Enns E, Marinier E, Mandal A, Herman EK, Chen C-Y, et al. Proksee: in-depth characterization and visualization of bacterial genomes. Nucleic Acids Res. 2023;51: W484–W492.

51. Vernikos GS, Parkhill J. Interpolated variable order motifs for identification of horizontally acquired DNA: revisiting the Salmonella pathogenicity islands. Bioinformatics. 2006;22: 2196–2203.

52. Arkin AP, Cottingham RW, Henry CS, Harris NL, Stevens RL, Maslov S, et al. KBase: The United States department of energy systems biology knowledgebase. Nat Biotechnol. 2018;36: 566–569.

53. Stothard P, Grant JR, Van Domselaar G. Visualizing and comparing circular genomes using the CGView family of tools. Brief Bioinform. 2019;20: 1576–1582.

54. Parks DH, Imelfort M, Skennerton CT, Hugenholtz P, Tyson GW. CheckM: assessing the quality of microbial genomes recovered from isolates, single cells, and metagenomes. Genome Res. 2015;25: 1043–1055.

55. Altschul SF, Gish W, Miller W, Myers EW, Lipman DJ. Basic local alignment search tool. J Mol Biol. 1990;215: 403–410.

56. Hall JPJ, Botelho J, Cazares A, Baltrus DA. What makes a megaplasmid? Philos Trans R Soc Lond B Biol Sci. 2022;377: 20200472.

