## Supplemental Materials for "Completed genomes from *Variovorax* provide insight into genome diversification through horizontal gene transfer"

Supplemental Data file for Genomics Paper

**R script for Phylogenetic Tree visualization**

\# Install Libraries, BiocManger

if (!requireNamespace("ape", quietly = TRUE)) install.packages("ape")

if (!requireNamespace("phytools", quietly = TRUE)) install.packages("phytools")

#Libraries Required

library(ape)

library(phytools)

#Path and Directory Use

setwd("/Users/christopherneville/Desktop/Yeast_Mouse/Masters/")

file_path <- "Complete_Variovorax_Genomes_Tree_better-labels2.newick"

pdf("phylogenetic_tree_high_res3.pdf", width = 20, height = 14) #This starts plotting to this file, until the dev.off line, If using Rstudio it will not show up in the plots

#File read in's & Checks

phylo_tree <- read.tree(file_path)

print(phylo_tree)

\# Plotting using ape/Phytools

plot.phylo(phylo_tree, type = "phylogram", edge.width = 2, cex = 0.8) # Plots the Phylo. Tree

nodelabels(text = phylo_tree$node.label, # adds in bootstrapping values

adj = c(-0.5, -0.5),

frame = "none",

cex = 0.8,

col = "black")

dev.off()

**Supplemental Table 1. Assembled Genomes and Secondary Replicons from NCBI as of 5/15/24 including contributions from this work.**

| **Strain** | **Replicons**  **(Mbp)** | **%(G+C)**  **each** | **Culture site** | **ref** |
| --- | --- | --- | --- | --- |
| *V. paradoxus* VAI-C | **6.67**  **2.48**  **0.29** | **69.5**  **68.5**  **60.5** | Univ. of Iowa | **MRA** |
| *V. paradoxus* EPS | **6.55** | **66.5** | CSUSB | **MRA** |
| *V. paradoxus* CSUSB | **5.57** | **65.5** | CSUSB | **MRA** |
| *V. paradoxus* 4MFCol3.1 | **5.42**  **1.60** | **67.8**  **66.7** | UNC |  |
| *V. paradoxus* 110B | **6.37**  **1.18** | **67.9**  **66.9** | UNC |  |
| *Variovorax* sp. 160MFSha2.1 | **6.78**  **0.57** | **67.9**  **64.8** | UNC |  |
| Variovorax sp. 278MFTsu5.1 | **6.86**  **0.47** | **67.9**  **65.0** | UNC |  |
| *V. paradoxus* 295MFChir4.1 | **5.56**  **1.63** | **67.7**  **66.7** | UNC |  |
| *V. paradoxus* 349MFTsu5.1 | **5.52**  **1.63** | **67.7**  **66.7** | UNC | **Identical second rep** |
| *Variovorax* sp. 350MFTsu5.1 | **6.87**  **0.56** | **67.9**  **65.1** | UNC |  |
| *V. paradoxus* 369MFTsu5.1 | **5.56**  **1.63** | **67.7**  **66.7** | UNC | **Nearly identical to 295/349** |
| *Variovorax* sp. 375MFSha3.1 | **6.89**  **0.46** | **67.8**  **65.1** | UNC | **Finkel et al** |
| *V. paradoxus* S110 | **5.63**  **1.13** | **67.6**  **67.0** | RPI (potato)  AHL degrader | **Han et al** |
| *V. paradoxus* ATCC17713 | **5.52**  **1.15** | **67.8**  **67.4** |  |  |
| *Variovorax* sp. KK3 | **6.99**  **0.26** | **67.6**  **64.3** | Georgia Tech | **KK cite** |
| *Variovorax* sp. NFACC26 | **7.75**  **0.55** | **67.5**  **65.0** | **Noble** |  |
| *Variovorax* sp. NFACC27 | **8.32** | **67.4** | **Noble** | **Integrated 2^nd^ rep** |
| *Variovorax* sp. NFACC28 | **7.73**  **0.55** | **67.5**  **65.0** | **Noble** |  |
| *Variovorax* sp. NFACC29 | **7.77**  **0.55** | **67.5**  **65.0** | **Noble** |  |
| *Variovorax* sp. PBL-H6 | **5.99**  **0.84**  **0.042** | **67.0**  **62.5**  **63.5** | Linuron contaminated soil, Belgium | **Ozturk et al** |
| *Variovorax* sp. PBL-H4 | **6.43**  **0.12**  **0.10** | **67.0**  **65.0**  **62.5** | Linuron contaminated soil, Belgium | **Ozturk et al** |
| *Variovorax* sp. PBL-E5 | **5.66**  **0.80**  **0.55**  **0.071** | **67.5**  **62.5**  **67.0**  **61.5** | Linuron contaminated soil, Denmark | **Ozturk et al** |
| *Variovorax* sp. PMC12 | **5.87**  **1.14** | **67.5**  **67.5** | Potting soil,  Wanju SK |  |
| *Variovorax* sp. PAMC26660 | **7.4** | **66.0** | Antarctic soil |  |
| *Variovorax* sp. PAMC28562 | **4.7** | **63.5** | Antarctica |  |
| *Variovorax* sp. PAMC28711 | **4.3** | **66.0** | Himantormia lichen, Antarctica |  |
| *Variovorax* sp. RKNM96 | **7.17** | **66.5** | Kamloops, Battle Bluff, British Columbia |  |
| *Variovorax* sp. WDL-1 | **6.72**  **0.82**  **0.57**  **0.21**  **0.025 0.020** | **67.0**  **62.5**  **63.5**  **63.5**  **62.5**  **62.5** | Linuron contaminated soil, Belgium | **Ozturk et al** |
| *Variovorax* sp. 38R | **6.87** | **67.5** | France, atrazine deg | **MRA**  **2021** |
| *Variovorax* sp.SRS16 | **5.76**  **0.80**  **0.56**  **0.48**  **0.071** | **67.5 62.5**  **64.5**  **67.0**  **61.5** | Linuron contaminated soil, Denmark | **Ozturk et al** |
| *Variovorax* sp. PDNC026 | **5.87**  **1.24** | **67.5**  **67.5** | Plastic debris, St Paul, MN |  |
| *Variovorax* sp. RA8 | **6.50**  **0.43**  **0.43**  **0.068** | **67.0**  **65.0**  **64.0**  **61.0** | Linuron contaminated soil, Belgium | **Ozturk et al** |
| *V. paradoxus* 2u118 | **5.62**  **1.30** | **67.5**  **67.0** | Legume root  UCLA |  |
| *V. paradoxus* 5C-2 | **7.29** | **67.5** | Russia | **Cite** |
| *V. paradoxus* B4 | **5.80**  **1.35** | **67.0**  **67.0** | Goettingen Genomics Laboratory |  |
| *V. paradoxus* JBCE486 | **7.29** | **66.0** | Jeonbuk National University, South Korea |  |
| *V. paradoxus* MGMM5 | **5.81** | **65.5** | Russia |  |
| *V. paradoxus* SPNA7 | **5.88**  **1.19** | **67.5**  **67.5** | UCLA Legume root | **NCBI** |
| *V. boronicumulans* EBFNA2 | **6.46**  **0.98** | **67.5**  **65.0** | UCLA  Legume root |  |
| *V. boronicumulans* J1 | **7.14** | **68.0** | China |  |

| 1. Supplemental Table 2. Core genes in secondary replicons 2. Strain 2° replicon Core gene in 2° replicon | | |
| --- | --- | --- |
| 4MFCol3.1 | Chromid | *aro*B, *pyr*F, *lsp*A |
| 110B | Chromid | None |
| 160MFSha2.1 | Megaplasmid | *lsp*A |
| 278MFTsu5.1 | Megaplasmid | *omp*H, *lsp*A |
| 295MFChir4.1 | Chromid | *aro*B, *lsp*A |
| 349MFTsu5.1 | Chromid | *aro*B, lspA |
| 350MFTsu5.1 | Megaplasmid | None |
| 369MFTsu5.1 | Chromid | *aro*B, *lsp*A |
| 375MFSha3.1 | Megaplasmid | *omp*H, *lsp*A |
| VAI-C | Chromid and  Plasmid | Chromid: *his*D , *mlt*A , *lsp*A  Plasmid: *rmu*C |
| ATCC17713 | Chromid | *lsp*A |
| KK3 | Plasmid | *hem*C |
| NFACC26 | Megaplasmid | *his*D |
| NFACC28 | Megaplasmid | *his*D |
| NFACC29 | Megaplasmid | *his*D |

*aro*B, dehydroquinate synthase

*pyr*F, orotidine 5’ phosphate decarboxylase

*lsp*A, peptidase A8 (signal peptidase)

*omp*H, porin

*his*D, histidinol dehydrogenase

*mlt*A, membrane bound lytic murein transglycosylase A

*rmu*C, recombination limiting protein C

*hem*C, porphobilinogen deaminase

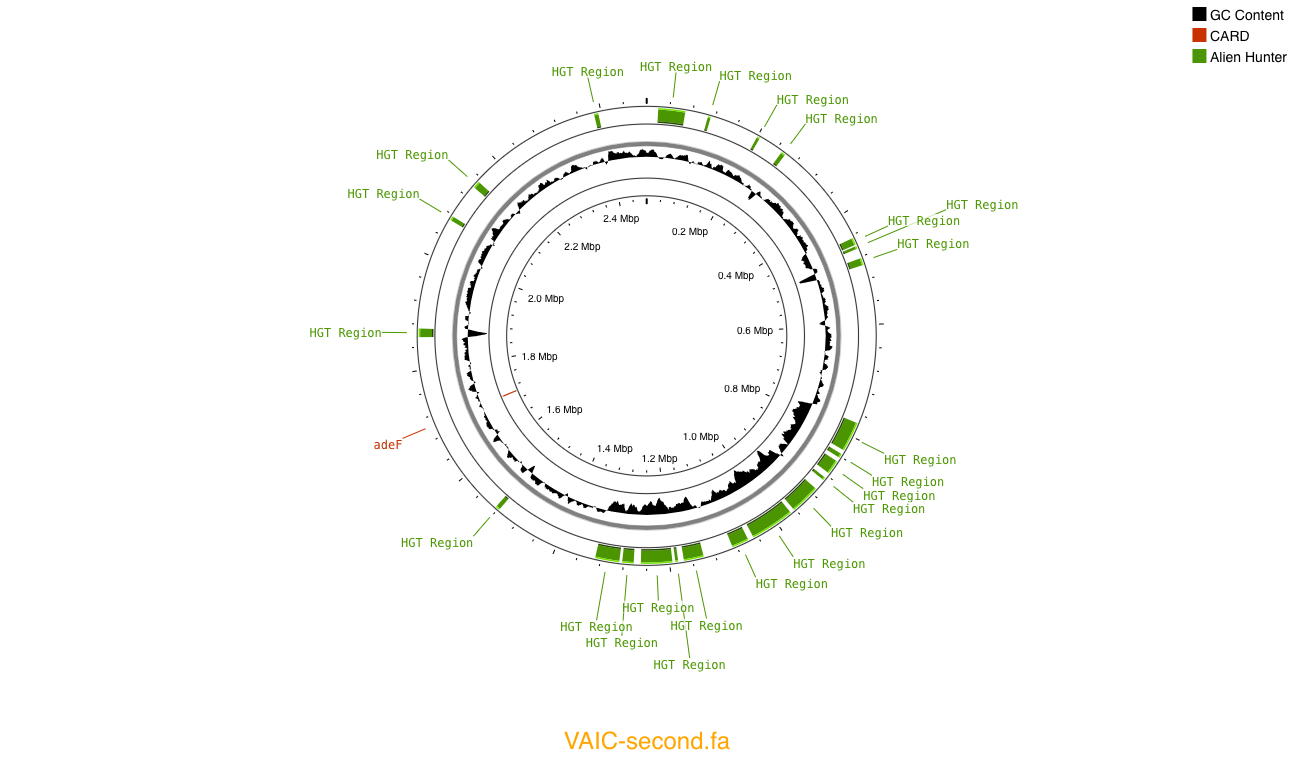
**Supplemental Figure 1. VAI-C chromid map constructed in Proksee (**[**www.proksee.ca**](http://www.proksee.ca)**) showing large region of insertions based on G+C content and the Alien Hunter algorithm.**

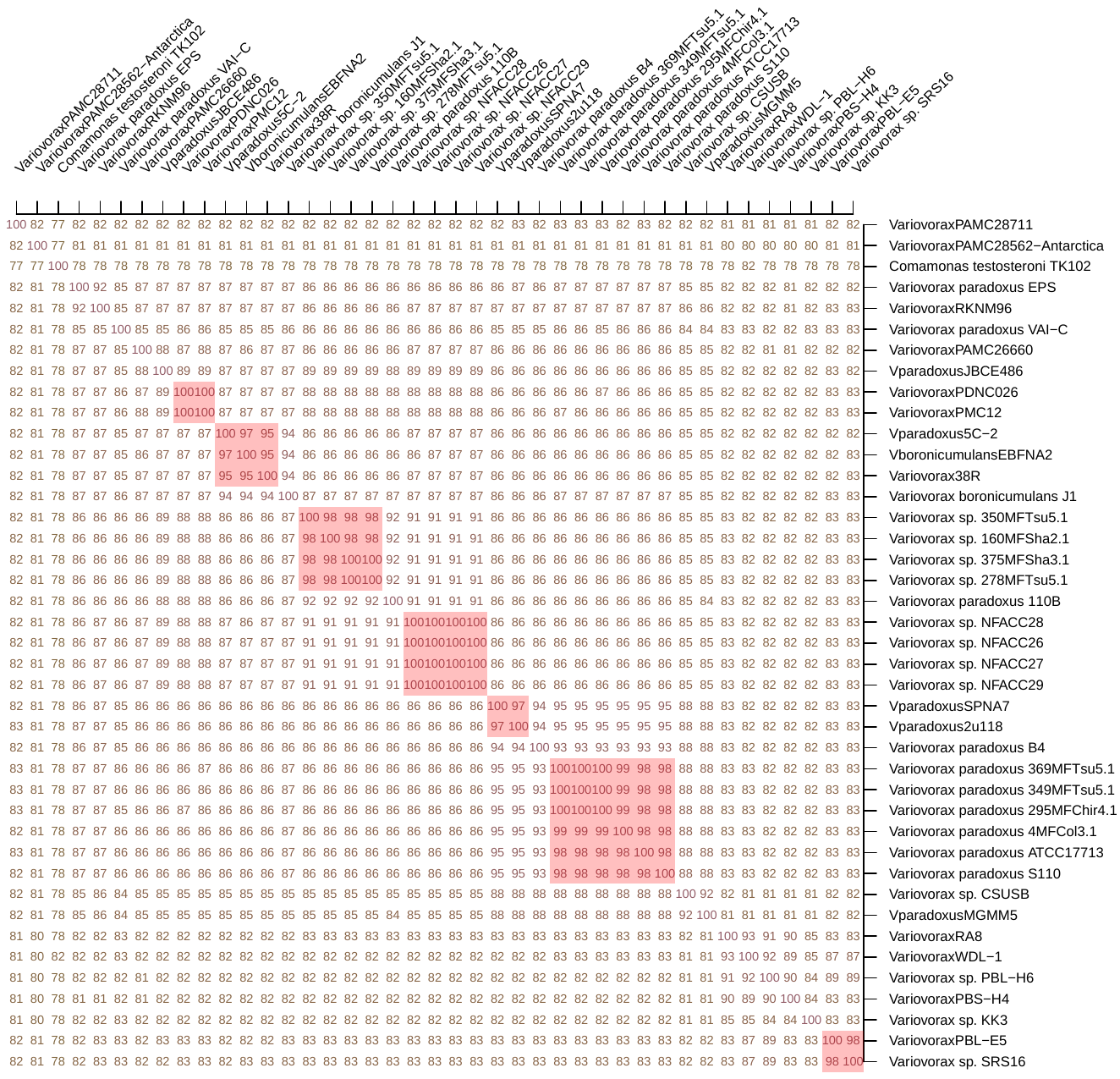

**Supplemental Figure 1. ANI Cluster Matrix of all completed *Variovorax* genomes**

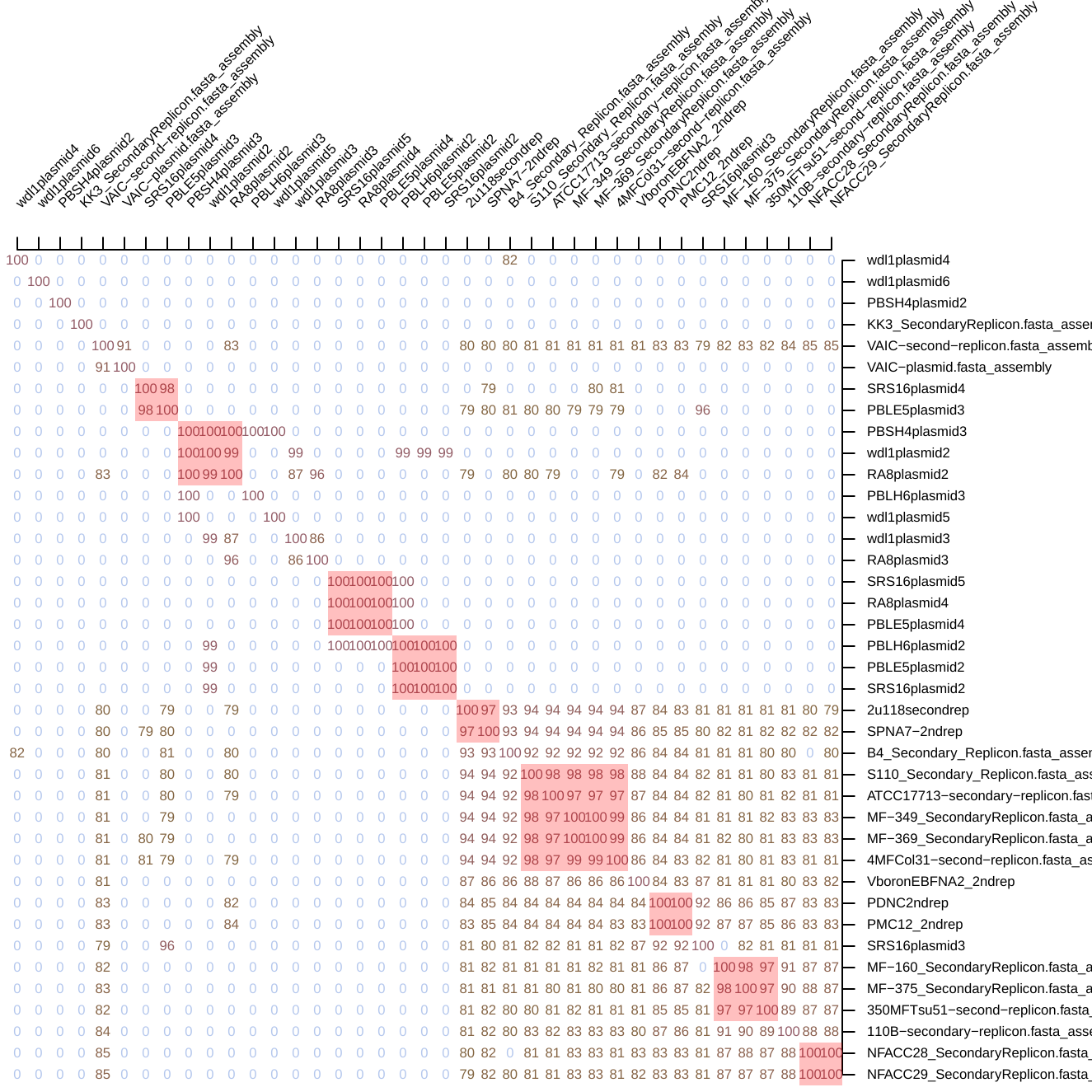

**Supplemental Figure 2 . ANI matrix for secondary replicons**
